# Evolutionary radiation of *Polaromonas* from mountain glaciers downstream

**DOI:** 10.64898/2026.02.18.706520

**Authors:** Grégoire Michoud, Aileen Geers, Hannes Peter, Amy C. Thorpe, Zhi-Ping Zhong, Virginia Rich, Tom J. Battin

## Abstract

Habitat transitions are central to microbial ecology and evolution and have been extensively studied across vastly different environments, such as between saline and non-saline environments. However, microbial habitat transitions along other large-scale environmental gradients remain poorly studied. This is particularly true for transitions involving the cryosphere, despite building evidence suggesting the Cryogenian as important for evolutionary radiation. Here, we investigated ecosystem transitions and related genomic adaptations of the cosmopolitan cryospheric *Polaromonas* bacterium. We constructed a pangenome from 282 high-quality genomes, sourced from glaciers, glacier-fed streams, lakes, wetlands, groundwater, rivers, and soils. Phylogenetic reconciliation revealed that the ancestral *Polaromonas* genome radiated from glacier ecosystems into various downstream environments through multiple independent transitions. These transitions were marked by extensive horizontal gene transfer and gene loss, with mobile genetic elements, such as plasmids and prophages playing key roles in genomic diversification. Predicted ancestral genomes encoded versatile metabolic and stress-response capacities, supporting adaptation to fluctuating and extreme conditions in the various cryospheric habitats. Compared to the ancestral *Polaromonas* genome, distinct genomic signatures were associated with specific habitats: glacier-fed stream lineages possess expanded stress tolerance repertoires, glacier lineages gained chemolithotrophic and anaerobic pathways, lake and wetland genomes acquired phototrophic functions, and soil lineages expanded substrate transport and stress tolerance. Together, our findings highlight the role of genomic plasticity in the ecological success of *Polaromonas*, and also underscore the cryosphere as a potential evolutionary cradle from which lineages dispersed and adapted to downstream aquatic and terrestrial environments.

## Introduction

Ecological or habitat transitions, such as the conquest of land by plants or transitions between marine and freshwater environments, are a major driver of evolution across the tree of life^1–4^. Plant terrestrialization, which necessitated genomic adaptations to high irradiance, desiccation and oxygen stress^5^, represents a paragon among habitat transitions. Recent evidence points to the Cryogenian period (i.e. Snowball Earth) as potentially important for plant terrestrialization and evolution of multicellularity^6,7^. In contrast to eukaryotes, records of habitat transitions and their importance for the ecology and evolution of prokaryotes focused on few events such the marine–freshwater divide, soil and aquatic ecosystems, host-associated lifestyles, or endosymbiotic events^2,8,9^. Habitat transitions of bacteria are often associated with functional adaptations mediated by genome streamlining and horizontal gene transfer^2^. For instance, the transition of members of *Methylophilaceae* from freshwater sediments to ocean pelagic habitats^10^ or, the transition from marine to freshwater environments of SAR11^11^. In general, genes related to stress response, motility, energy and carbon metabolism are often associated with bacterial habitat transitions^2^. Despite their important ramifications for bacterial ecology and evolution, how genomic underpinnings of habitat transitions are related to the present-day success of bacterial clades remains generally understudied.

High dispersal capacity, metabolic versatility, stress tolerance, the ability to take up exogenous DNA and a large genomic pool allowing for adaptation are key requirements for successful habitat transitions^2^. Moreover, cryospheric environments, characterized by high spatio-temporal heterogeneity and connectivity between frozen and liquid habitats may represent hotspots for habitat transitions — as exemplified by plant terrestrialization^7^. Similar to global glaciation during Snowball Earth, the cryosphere continues to be a highly selective environment for microbial life^12^. Microbial clades specialized to the cryospheric conditions are adapted to cope with low temperatures, osmotic and radiative stress, and ultra-oligotrophy^12^.

*Polaromonas* is an iconic bacterial genus with a cosmopolitan distribution and reported from numerous cryospheric (e.g., glaciers, permafrost soils)^13–16^ and non-cryospheric (e.g., wetlands, groundwater) ecosystems^17–19^. *Polaromonas* is metabolically versatile^20^, possessing genes involved in both heterotrophic processes able to degrade organic carbon^21^ and genes involved in the degradation of recalcitrant organic compounds such as hydrogen^22^ or arsenic^23^. Some species are even able to fix nitrogen^24^ although this metabolic pathway is not widely distributed within the genus^20^. Further, in contrast to *Prochlorococcus*, a cosmopolitan marine bacterium^25^, the *Polaromonas* genome is not streamlined; it has an average size of 3 to 5 Mb and a GC content of 50-60%^21,26^, typical for cryospheric taxa^12^. Comparative genomics has revealed the role of horizontal gene transfer in shaping the genetic diversity and adaptive potential of *Polaromonas* to cold environments^27^. Compared to mesophilic species (e.g. *Polaromonas naphthalenivorans* CJ2), cold adapted *Polaromonas* possesses more (approximately 10%) horizontally transferred and transposase genes, collectively suggesting a greater potential for genome rearrangement and acquisition of novel genetic material^27^.

The apparent ease at which *Polaromonas* disperses over long distances^20,28^ may facilitate habitat transitions and explain its occurrence in air and snow at high altitudes^29–31^. Furthermore, local selection pressures and interactions of *Polaromonas* with other bacteria and also algae, drive niche separation and microevolution within the genus^32^. Such microevolution may lead to the formation of *Polaromonas* phylotypes associated with distinct environments as reported from glacier habitats^33^, and further highlighting the adaptive capabilities of the genus. Indeed, a recent study has shown a clear functional selection between eight *Polaromonas* found in arctic environments compared to those found elsewhere (e.g. glaciers, groundwater and wastewater), where the former tended to have extended genomes (on average of 500 kb) with an increase of genes related to osmoprotection and carbohydrate metabolism^34^. This study however did not address the transition and evolutionary history of *Polaromonas* across cryospheric environments and from these to non-cryospheric ones. One hypothesis related to this radiation across environments is related to ecosystem relocation on snowball earth^35^, where the expansion of polar–alpine biomes across earth allowed the migration of adapted bacteria allowing them to survive and evolve and later repopulate post-glacial environments. The Cryogenian Snowball Earth, 650 million years ago, was important for evolution and radiation of various lineages, such as Cyanobacteria or Chlorophytes^35^, and through habitat transitions into various environments that emerged upon deglaciation. The Cryogenian also left genomic and physiological imprints on bacteria, such as *Prochlorococcus* which is a widespread primary producer in the modern ocean^36^. It seems therefore intuitive to consider glaciers as cradles of bacterial diversity, facilitated by habitat transitions as meltwaters and contained cells flow through rivers and lakes, and into terrestrial ecosystems during inundations. Further, glaciers can also be considered as “refugia” for cryospheric microbes during periods of deglaciation, thus providing longer-term cold habitat^37^. Therefore, we argue that *Polaromonas* is a good model system to study bacterial habitat transitions, even the potential role of the cryosphere for its evolution.

In this study, we explored the *Polaromonas* pangenome containing 282 high-quality genomes sourced from a wide range of cryospheric (e.g., glaciers, permafrost soils) and non-cryospheric (e.g., wetlands, lowland rivers, groundwater) ecosystems. This enabled us to explore the breadth of genomic variation across *Polaromonas* populations and unravel mechanisms that allow these bacteria to colonize diverse and often extreme environments. Using phylogenetic reconciliation^38^, we provide novel insights into genome evolution of *Polaromonas*, assess its ancestral state and rearrangement of its genetic repertoire as it transitioned across habitats. Collectively, our findings shed new light on the ecology and evolution of a most successful bacterial clade, with its roots likely in the world’s glaciers and their streams that are now changing at a rapid pace because of climate change. Our results support the notion of the cryosphere as playing a role in the microbial diversification with impacts on downstream ecosystems. Therefore, our findings provide a microbial perspective on how the cryosphere has shaped and continues to influence the global genetic and ecological landscape.

## Results and discussion

### Distribution and diversity of *Polaromonas*

First, we explored the spatial distribution of *Polaromonas* to assess its capacity to colonize and dwell in various environments. To this end, we analysed 16S rRNA gene data from the Microbe Atlas Project^39^. We found *Polaromonas* to be present across all continents and in a wide range of ecosystems and habitats, including rivers, lakes and reservoirs, groundwater, glacier ice and cryoconite, soils, sediments, and both animal and plant hosts (Figure S1). The majority of *Polaromonas* records were located in aquatic environments, including glaciers and ice (243, 63.4%), followed by soils and sediments (126, 32.9%), and animal and plant hosts (14, 3.65%). In ocean or saline lakes, *Polaromonas* was found only at low relative abundances (< 0.1% relative abundance), with the notable exception of estuaries with low temperature and low salinity (e.g. Finland and Antarctica)^28,40^, suggesting that the genus has not globally passed the salinity barrier^41^.

In a next step, we compiled a list of genomes to investigate the pangenome of *Polaromonas*, including both MAGs, SAGs and genomes sequenced from isolates. We collected genomes from the NCBI genbank^42^, Tibetan aquatic microbiomes^43^, Tibetan glacier microbiomes^44^, glacier-fed streams microbiomes (GFS)^16^, Guliya ice-cap^45^, UK^19^ and US river microbiomes^18^. These genomes were filtered for quality (*Methods*) and taxonomies were confirmed using the genome taxonomy database (GTDB)^46^. We retained 282 non-redundant genomes from a wide range of cryospheric and non-cryospheric environments, including glaciers (73), GFS (54), lowland rivers (17), wetlands (26), lakes (55), groundwater (23), soils (31), and other habitats (e.g. where isolates were found)(3) (Table S1, Table S2 and Figure 1a-b).

**Figure 1:**
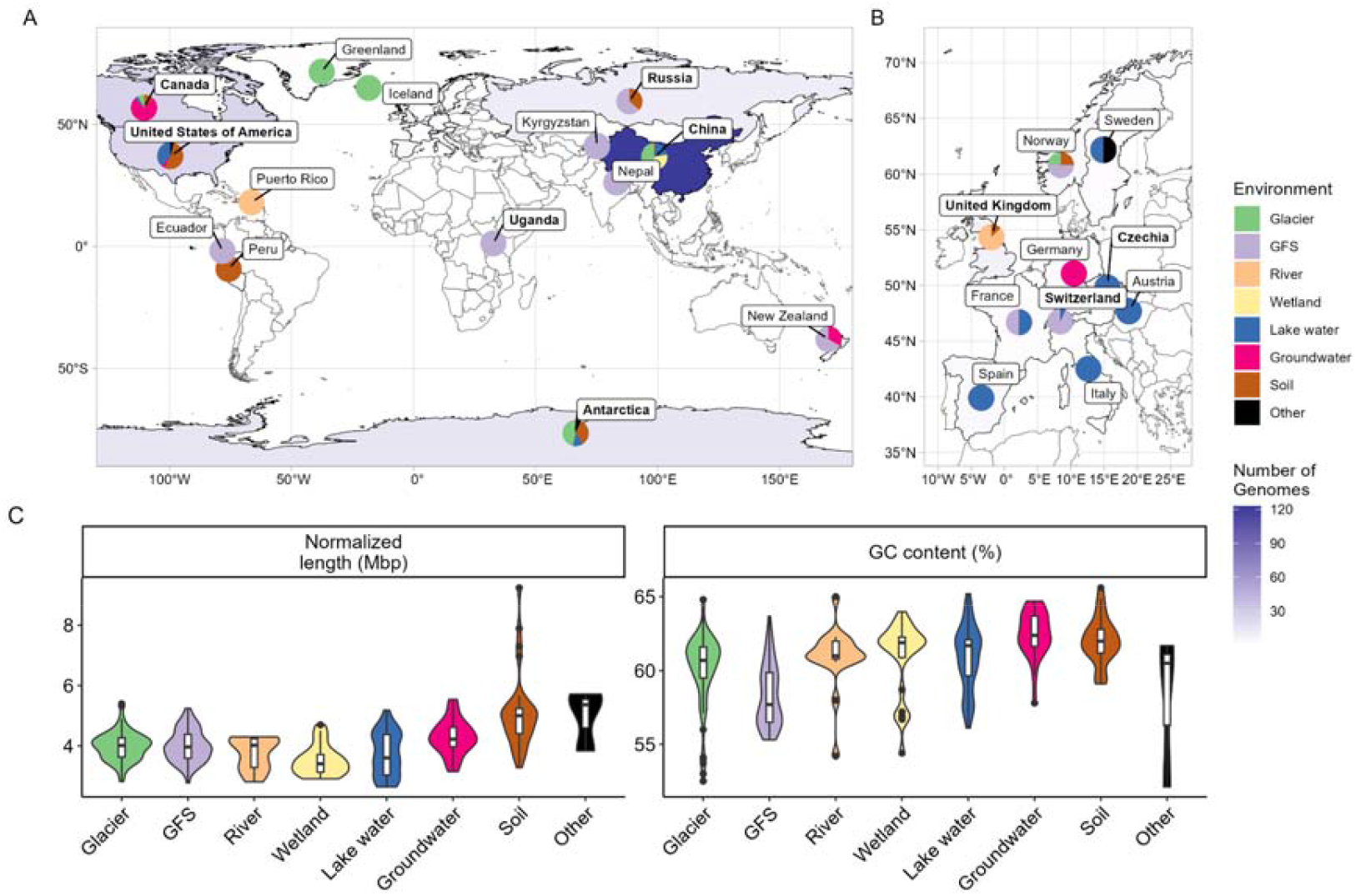
(a, b) Locations of the origins of *Polaromonas* used in this study. For each country or region, the shading corresponds to the number of genomes obtained and the pie chart represents the environment from which these genomes were obtained. Panel b represents European countries for visualization purposes. The “other” category corresponds to the cultured isolates obtained from other environments. (c) Normalized length (by completeness) and GC content (%) of *Polaromonas* genomes across different environments.

Accounting for contamination (86.1 ± 9.9%) and completeness (1.4 ± 1.8%) *Polaromonas* genome length averaged 4.04 ± 0.8 Mbp, with an average GC content of 60.4 ± 2.5% (Table S2 and Figure 1c). *Polaromonas* genomes differed in size between environments with genomes sourced from soils were significantly larger (5.1 ± 1.26 Mbp (mean ± SD), Mann-Whitney U test, p < 0.05) than from the other environments, with the exception of groundwater genomes and type strains, which had similar sizes (Table S2 and Figure 1c). These results were expected as “terrestrial” genomes tend to be larger than “aquatic” genomes, and genomes from isolates are larger than MAGs^47^. The distribution of GC content was similar as previously reported and did not significantly differ between environments^34^.

### Genomic differentiation across environments

To explore how *Polaromonas* genomes differed across the various environments, we analysed the pangenome based on persistent, shell and cloud genes (*Methods*). We sourced a total of 958,785 genes from the 282 genomes, which we arranged into 214,664 gene families, further divided into 1,905 persistent, 11,772 shell and 200,987 cloud unique gene families (Figure S2). On average, each persistent, shell and cloud gene family was found in 61.3 ± 22.7%, 7.6 ± 4.9% and 0.6 ± 0.6% of the genomes, respectively (Figure S3). Based on the presence or absence of persistent genes, we observed a clear environmental segregation (Figure 2). This environmental segregation was consistently observed regardless of which gene categories were analyzed, but it was most evident for cloud (ANOSIM, R = 0.47, p = 0.001) and shell genes (ANOSIM, R = 0.51, p = 0.001), compared to persistent genes (ANOSIM, R = 0.35, p = 0.001) (Figure S4). This pattern can be attributed to the fact that cloud and shell genes are less frequently shared across the pangenome^48^ and are therefore likely less essential to the core functionality of the organism than persistent genes^49^. Hence they are more likely to encode functions that facilitate adaptation to specific environmental conditions, thus driving the clearer segregation across environments.

**Figure 2:**
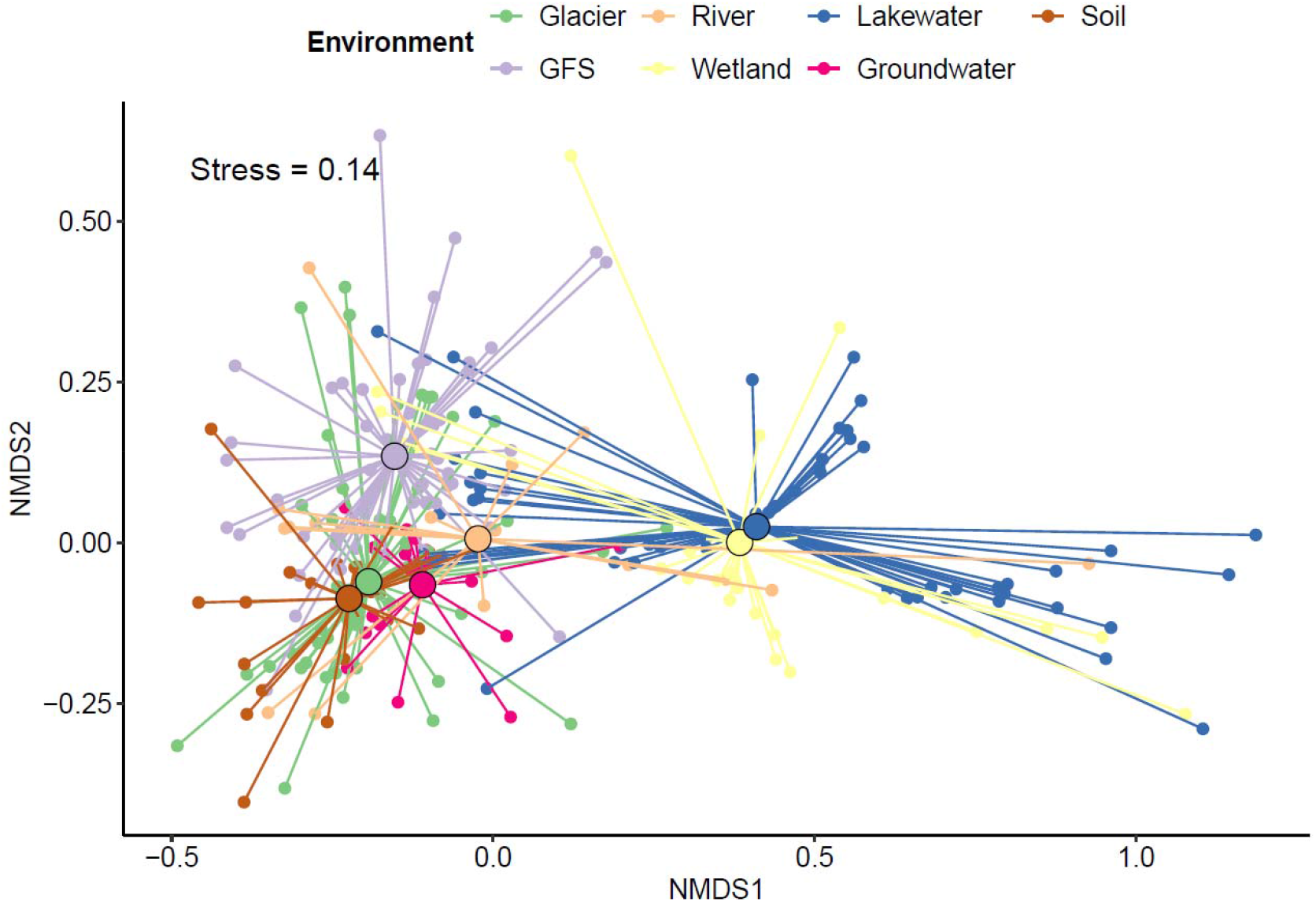
Non-metric multidimensional scaling ordination of persistent genes presence in different genomes show that *Polaromonas* cluster by source environment (analysis of similarities (ANOSIM), R = 0.35, P = 0.001).

A similar environmental segregation was previously observed based on a pangenome including 34 *Polaromonas* genomes with lower environmental resolution^50^ and a study comparing polar to non-polar *Polaromonas*^34^. Interestingly, we found clear clustering of genomes from lake water and wetlands on the one hand, and from GFS and glaciers, soils, lowland rivers and groundwater on the other (Figure 2). This suggests adaptations of *Polaromonas* strains to the different environments characterized by differences of gene repertoire. Similar findings have been reported for transitions of Planctomycetes from soil to freshwater and *Methylophilaceae* from lakes to oceans^10,51^. In those cases, many changes occurred through gene loss via genome streamlining to adapt to mostly oligotrophic conditions. Here, however, we do not observe signs of genome reductions, indicating different evolution modes to adapt to the various environments.

### Ancestral habitat and transitions

To further explore the importance of habitat transitions for the radiation of *Polaromonas*, we built a phylogenetic tree based on persistent and shell genes present in at least 30 genomes (i.e., ∼10% of the pangenome) (*Methods*). Our phylogenetic analysis revealed four main clusters, grouping most genomes from groundwater, lake water and wetland, GFS, and both soils and glaciers, respectively (Figure 3). This pattern suggests notable phylogenetic conservatism in habitat preference and consistent grouping of closely related lineages to similar environments. Hence, environmental filtering rather than dispersal would play a major role in shaping microbial compositions. Indeed, if dispersal were more frequent and successful, we would expect “mixed” phylogenetic signatures where related lineages would be found across multiple environments without clear clustering. Instead, the environmental clustering indicates that selective pressures are sufficiently strong to maintain habitat-specific lineages, limiting successful establishment outside their environments. These results would support the niche conservatism hypothesis stating that species tend to retain their ancestral ecological traits over time^52^. Furthermore, our results indicate that lineages often remain specialized to the environments occupied by their ancestors. As a result, closely related genomes are more likely to be found in similar habitats, while their transition across habitats seems rare.

**Figure 3:**
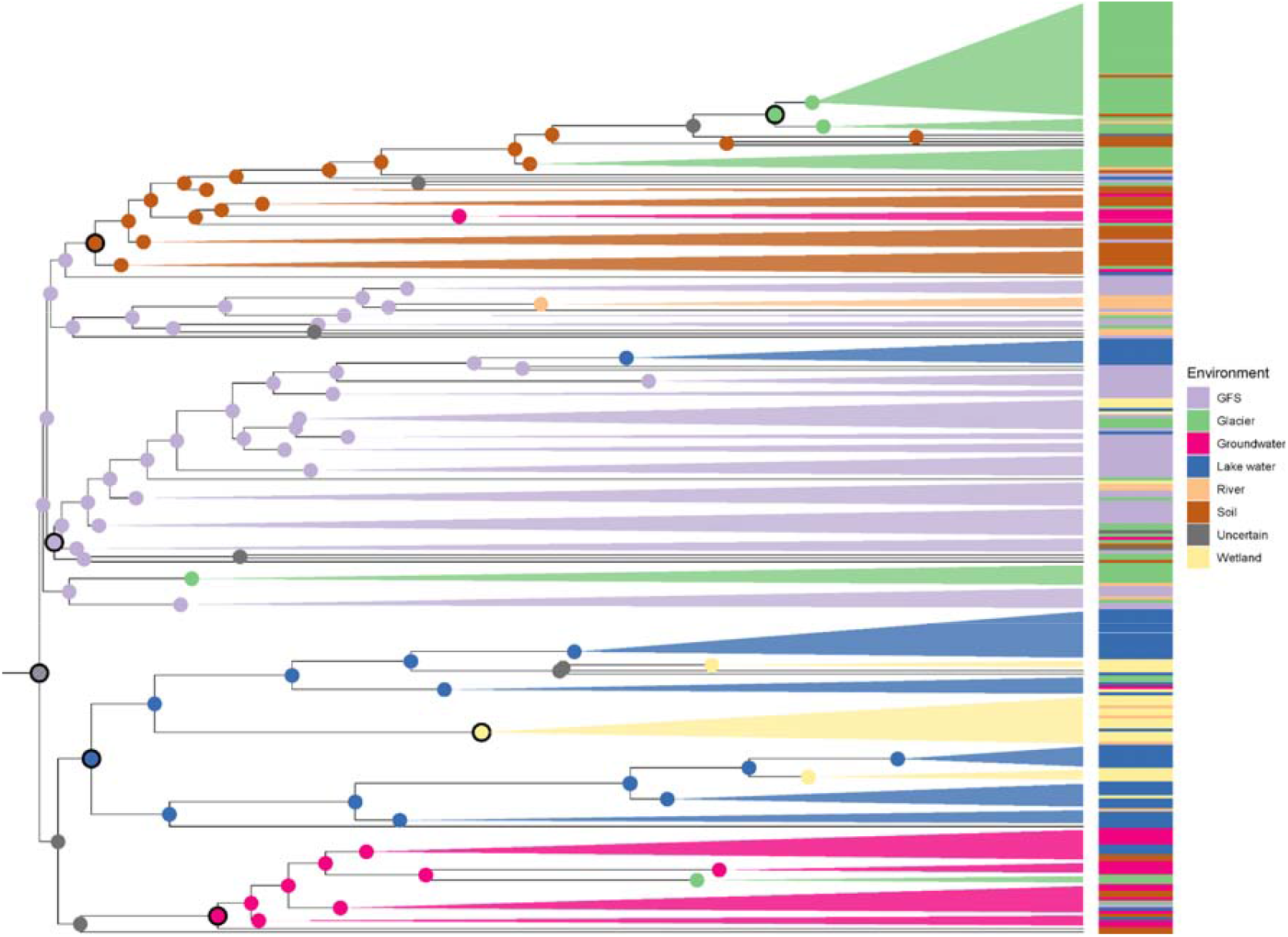
Phylogenetic tree of *Polaromonas* genomes. The color at each tip corresponds to the environment where the individual genomes were obtained, while the color at each node highlights the predicted environment where the *Polaromonas* ancestor was found via marginal posterior probabilities approximation using the pastML software^53^. Clade merging was also carried out using PastML, employing vertical merging, which groups adjacent sections of the tree where no state changes occur, and horizontal merging, which clusters independent events of the same type. Grey colors at the nodes indicate that there is uncertainty about the predicted environments. Highlighted nodes (in black) indicate predicted major habitat transitions.

Next, using pastML^53^, we predicted the potential ancestral habitat at each internal node of the phylogenetic tree and inferred habitat transitions of *Polaromonas* (*Methods*). Interestingly, pastML rooted the *Polaromonas* phylogenetic tree in GFS (Figure 3). We interpret this finding cautiously because predictions close to the root of the tree are particularly uncertain^54^.

Our analyses further predicted that *Polaromonas* transitioned from its putative ancestral environment in GFS into groundwater, lakes and wetlands, as well as soils and glaciers. The *Polaromonas* last common ancestor (LCA) has remained in GFS since an early ancestral stage, while other lineages have radiated to soils and then to glaciers. Another lineage diverged into groundwater, while a last one transitioned from lakes into wetlands.

While these broad patterns of habitat transitions appear relatively clear, we also detected signatures of multiple transitions occurring throughout the *Polaromonas* tree (Figure 3). Up to 10 MAGs from the same environments (e.g., rivers) were found throughout the phylogenetic tree, suggesting that some habitat transitions occurred several times. Numerous habitat transitions have been predicted to occur multiple times, which seems to be particularly true for host-associated bacteria, where some bacteria transition from their host to the environment very often, sometimes daily (e.g., *Vibrio fischeri* in a squid)^2,55^.

Furthermore, we hypothesize that transitions are linked to cycles of deglaciations during which ice sheets created “climate refugia”^37^ for cold adapted microbes. This was already suggested for some plants and animals especially in rock covered glaciers^56,57^, where climate refugia are areas large enough to support populations whose habitat is at risk due to changing climate. As glaciers retreated and expanded, dispersal strongly varied through hydrological mixing which may have facilitated genetic mixing and diversification of *Polaromonas* across some habitats. Noteworthy, we found genomes from lowland rivers throughout the tree and usually phylogenetically close to genomes from GFS in several small MAGs clusters (Figure 3). This indicates that lowland rivers act as collectors of *Polaromonas* MAGs from soil or groundwater, for instance^58,59^.

### Gene re-arrangements

To gain mechanistic insights in the gene re-arrangements that may accompany habitat transitions of *Polaromonas*, we focused on the internal nodes where the major habitat transitions are predicted to occur, besides the root (LCA). These transitions occurred at several stages, including the adaptation of LCA lineages to GFS, groundwater, lakes, and soils, as well as transitions of *Polaromonas* from lakes to wetlands and from soils to glaciers.

For each of these transitions, we determined the relative importance of duplications, transfers, losses and origination (i.e., DTLO) for each predicted gene family (Figure 3, Figure S5). While gene inheritance is usually vertical from mother to daughter cells, prokaryotic genes are susceptible to relatively high frequencies of transfers and losses, less so for duplications^38^. Briefly, transfers either occurring through horizontal gene transfers or genetic recombination tend to homogenize gene pools^60^. Gene loss is also frequent in prokaryotes and tends to reduce the metabolic capability of the cell in case of symbiosis but can also act as an adaptive evolutionary force useful when organisms face abrupt environmental challenges as encapsulated by the less-is-more hypothesis. This states that, for example, losing genes involved in the biosynthesis of compounds that are available in the environment, will allow the cells to use its energy for other purposes such as biofilms formation^61^.

We found that transfers (242.3 ± 149.4 per node) and losses (228.3 ± 160.1 per node) dominated DTLO events, compared to duplications (28.6 ± 42.7 per node) and originations (40.2 ± 107.3 per node). This pattern is consistent with general DTLO observations from prokaryotes (Figure S5)^38^.

As the number of horizontal transfers was relatively high, we further looked for genomic evidence of such events. First, we predicted the presence of plasmid and viruses in each genome using Genomad^62^. Due to the fact that most of the genomes were assembled from metagenomes, we acknowledge possible biases associated with this analysis. Nevertheless, we found that on average 95.2% ± 5.0% (mean ± SD) of the genomes possessed at least one plasmid, ranging from 80.6% to 100% of the genomes present in soils and rivers, respectively. Furthermore, many genomes either possessed a prophage or a virus (50.3% ± 14.5% of genomes); 80% of the wetland genomes are predicted to have been infected by viruses, while only 29% of glacier genomes were infected, potentially due to the higher cell densities in wetland or soils than glacier (Figure S6a). As expected most genes on these mobile genetic elements were predicted to have been transferred at the habitat transitions (Figure S6b). These results could explain why the *Polaromonas* genus is microdiverse, that is where adaptations are observed at a finer phylogenetic level than species^20^. Indeed the relevance of viruses for the diversity of gene pools across communities has been documented, and it has been suggested that the maintenance of high strain diversity prevents the accumulation of resources by specific strains^63,64^. The high abundance of mobile genetic elements and their presence at habitat transitions could facilitate the adaptation of *Polaromonas* species to changing environmental conditions, either by selecting against other populations or allowing selected species to diversify in the different environments.

### The last common ancestor of *Polaromonas*

Having unveiled major habitat transitions of *Polaromonas*, we further assessed the genomic traits of its LCA (*Methods*). First, we note that unsurprisingly, the vast majority of DTLO events were originations (315) (Figure S5). Our results suggest a range of physiological and metabolic traits that enabled the LCA to adapt to diverse and fluctuating environmental conditions (Figure 4, 5). Most likely, it displayed the capacity for aerobic respiration and heterotrophy (i.e., ETC complex IV) allowing for the exploitation of various organic energy sources. Furthermore, our predictions suggest that it was able to synthesize and export extracellular polymeric substances (EPS), commonly associated with biofilm formation. It also possessed genes related to pH (decarboxylase and antiporter systems) and temperature stress (heat-induced transcriptional regulators). Finally, the *Polaromonas* LCA likely possessed diverse substrate uptake systems, including transporters for carboxylates (both dicarboxylates and tricarboxylates), simple and complex carbohydrates (including a complete Entner-Doudoroff pathway), free amino acids, metal ions, nitrogenous compounds, organophosphorus compounds, and vitamin B.

**Figure 4:**
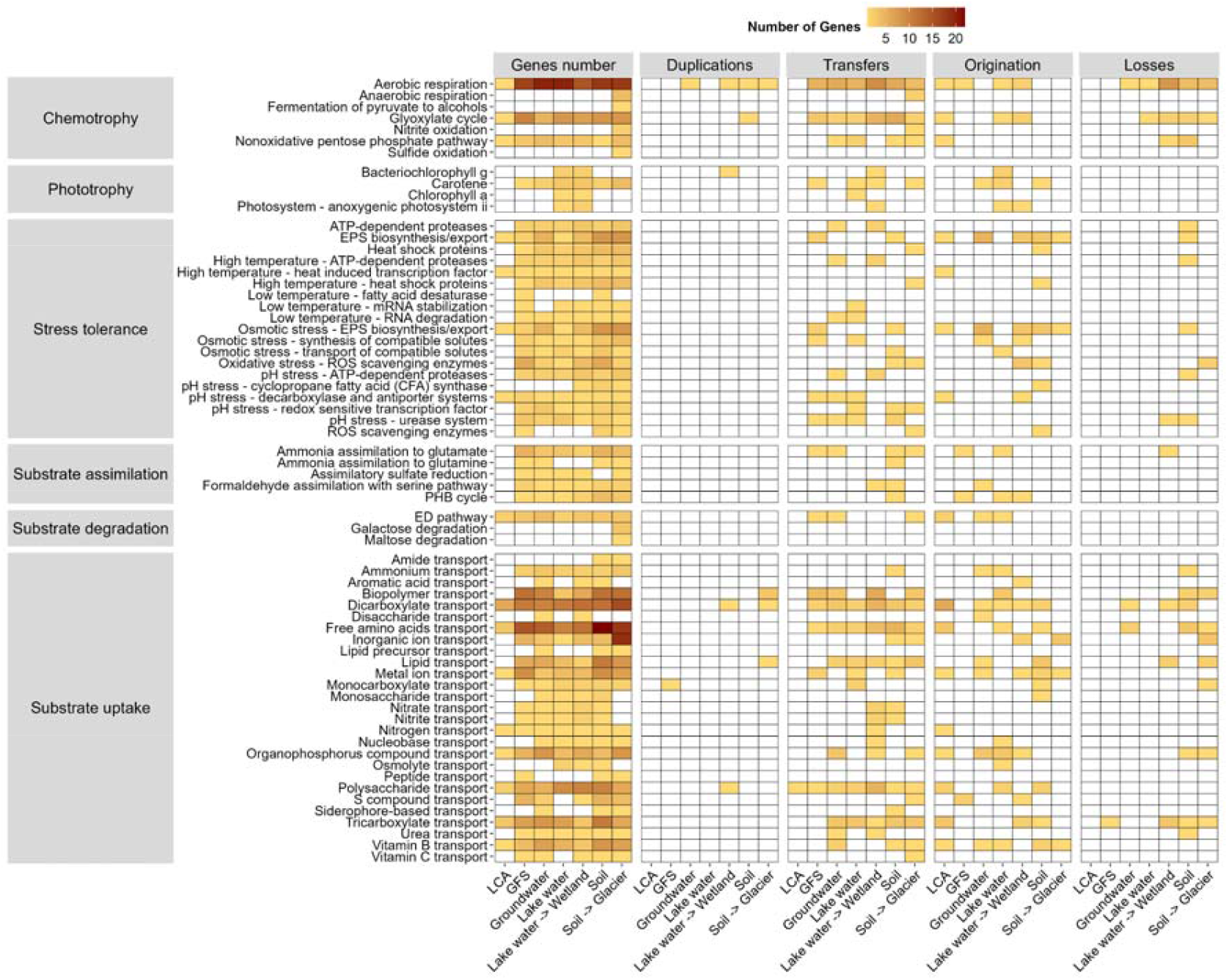
Heatmap representing duplications, transfers, originations, and losses of gene families associated with metabolic, stress response, and transport functions across six habitat adaptations and transitions of *Polaromonas*. Gene numbers correspond to numbers of representative genes per function (as determined by microtrait^66^) found in the different genomes, while color intensity reflects the number of genes per category.

**Figure 5:**
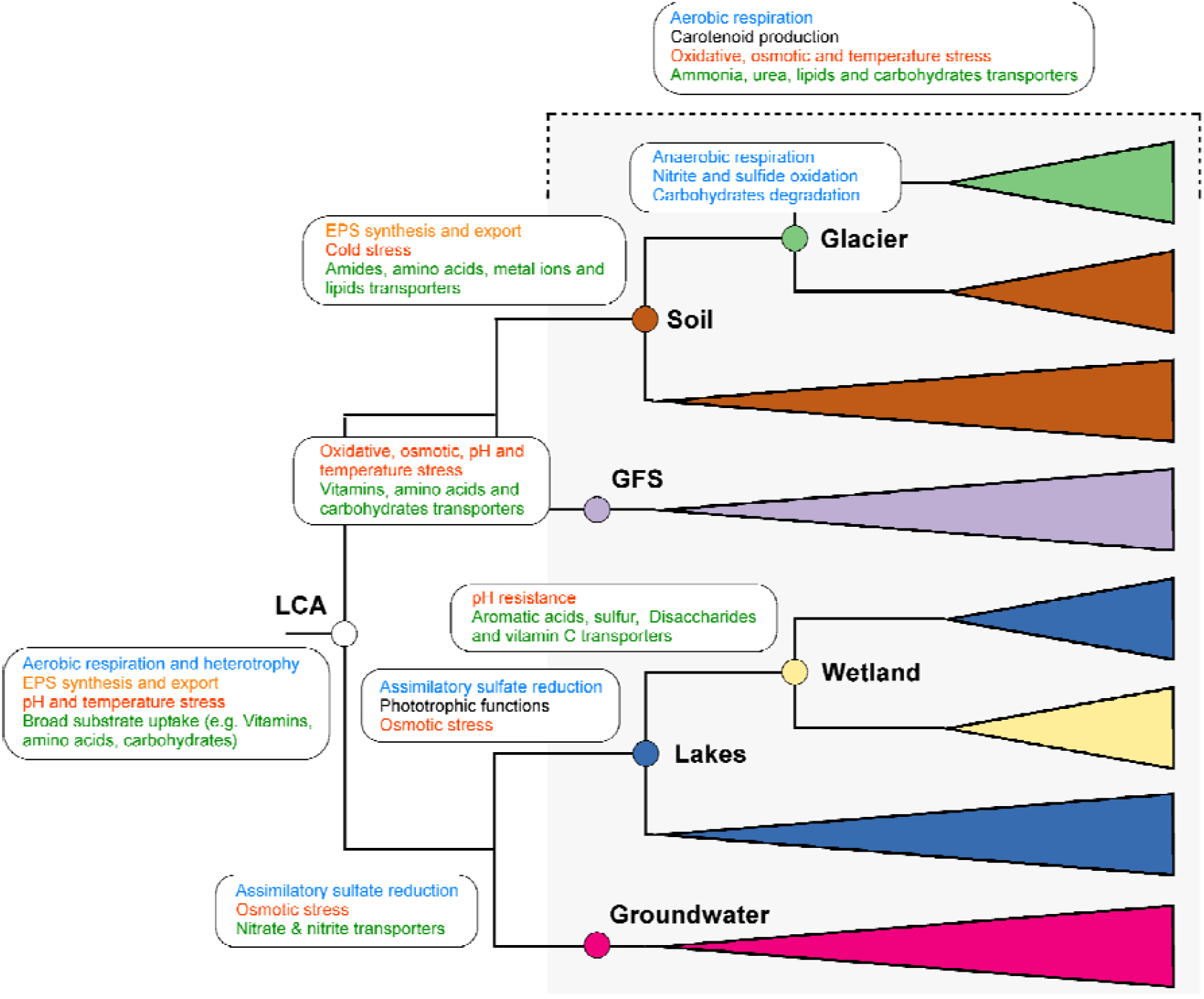
Schematic representation of functions gained amongst different habitats transitions. The grey square represent function gains that occurred in all habitat transitions

Together, these traits are consistent with a biofilm mode of life capable of exploiting a wide range of substrates. Indeed, these traits are advantageous to dwell GFS where biofilms reduce turbulent-induced erosion of microbial cells and energy sources are fluctuating in time^16,50,65^. Although environmental predictions suggest that the *Polaromonas* LCA dwelled in GFS, the relatively small number of genes predicted for its genome (330) compared to those at other habitat transitions (735.7 ± 219.8) makes functional comparisons challenging. In fact, the absence of a gene in these predicted genomes cannot be taken as definitive evidence of its true absence. Hence, we did not consider that the LCA transitioned directly from GFS to other ecosystems; rather, we considered adaptations of the LCA to specific environments (GFS, soils, lakes, and groundwater), as well as transitions from lakes to wetlands and from soils to glaciers.

### Genomic adaptations to habitat transitions

The traits of the *Polaromonas* LCA diversified with each transition downstream to aquatic and terrestrial ecosystems (Figure 4, 5). Overall, predicted node-specific genomes at the various habitat adaptation and transitions (GFS, groundwater, lakes, soils, transitions from lakes to wetland, and transitions from soil to glaciers) tended to possess more metabolic genes compared to the LCA genome. Genomes downstream from the LCA were characterized by a more complete aerobic respiration mechanism with the four clusters of electron□transport□chain complexes I–IV. Basic chemotrophy functions were also present in all the genomes at the habitat transitions, such as nonoxidative pentose phosphate pathway, the glyoxylate cycle and ammonia assimilation pathways (i.e., to glutamate). We also note the presence of genes involved in the biosynthesis of a carotenoid, neurosporene, an intermediate used in the biosynthesis of lycopene and other carotenoids, which are involved in the protection against UV radiation and oxidative stress^67^. In addition to the many transporters observed in the LCA genome, the genomes present at different habitat transitions possessed genes related to the substrate uptake of ammonium, urea, biopolymer, lipids, and polysaccharides. Furthermore, genes related to stress such as heat shock proteins, ATP-dependent proteases, transport of compatible solutes and reactive oxygen species (ROS) scavenging enzymes may afford protections against high temperature, elevated salt conditions and oxidative damage. Collectively, this gene repertoire allows *Polaromonas* species to exploit diverse carbon and nitrogen sources while withstanding thermal, osmotic, pH, and oxidative stresses. These traits facilitated the radiation of *Polaromonas* into the various aquatic and terrestrial ecosystems.

Despite the genetic repertoire that genomes at different habitat transitions shared, we note distinct genomic differentiations occurring depending on the environment where those genomes were found (Figure 4, 5). For example, genomes that transition to GFS possess numerous genes related to the adaptation to cold temperatures — surprisingly also elevated temperatures — including cold shock proteins, fatty acid desaturases, and RNA-stabilizing enzymes along with heat shock proteins. Furthermore, they also possess genes related to oxidative and osmotic stress, probably to counteract the effect of the high ultraviolet radiation at high altitudes, and redox shifts associated with periodic drying of GFS. An elevated number of genes related to the transport of amino acids, peptides, sugars, organophosphates, and vitamins collectively hint at interactions with algae and recycling of nutrients within biofilms in GFS^16,50,68^. Interestingly, a relatively high proportion of these genes were predicted to be horizontally transferred from other *Polaromonas* strains. We interpret this gene repertoire of *Polaromonas* as an adaptive response to the extreme and fluctuating environment (e.g., water flow, temperature and light) in GFS, driven by glacier melt dynamics^15,69^. We note that the environment of GFS is more extreme and less stable compared to wetlands, soils and groundwater, for instance.

Along the hydrological continuum, the transition into lakes from the *Polaromonas* ancestor was marked by several and strikingly important genomic modifications that facilitate a planktonic mode of life (Figure 4, 5). In fact, our results suggest that *Polaromonas* genomes are significantly enriched in phototrophic genes with the additions of functions related to the biosynthesis of both chlorophyll *a* and *g* as well as the biosynthesis of lycopene and an anoxygenic photosystem. These modifications mostly occurred through originations and gene transfers, and were not found in the other habitat transitions, highlighting the plasticity of *Polaromonas*. We further note that anoxygenic photosystem systems are oxygen-independent, ecologically widespread, and critical for microbial energy harvesting in anaerobic, low-light environments. Hence, they are well adapted to the pelagic environment in lakes and wetlands. We also observed the presence of osmolyte transporters (two genes with one origination), probably involved in salt stress regulation. Similar to the groundwater transition, genomes at the lake transition also possess genes related to assimilatory sulphate reduction. Predicting that wetland *Polaromonas* genomes descended from lake genomes, we observed the gain of different transporters for aromatic acid, disaccharides, vitamins C and sulphur compounds, thus extending its capacity for scavenging a wide array of organic molecules. The presence of sulfur transporters is not surprising as wetlands are known to harbour very high concentrations of organic and sulfur compounds^70^. Wetland genomes also gained genes involved in pH resistance such as cyclopropane fatty acid synthase for membrane stabilization. Overall, many of the genes that differed from those in lake water appear to have been horizontally transferred, suggesting adaptations to the fluctuating environment of wetlands (e.g., changing redox conditions linked to drying).

If as predicted, the *Polaromonas* LCA had dwelled in GFS, it appears intuitive that it colonized groundwater through the continuous exchange between these water sources. We found that the transition of the *Polaromonas* LCA to groundwater occurred with the expansion of transporters related to nitrogen, particularly nitrate and nitrite transporters. Similarly, more stress tolerance genes are present with low□temperature mRNA□degradation genes and explicit heat□induced transcription factor regulation. Further, similar to the lake transition, we found the potential for assimilatory sulphate reduction pathways, revealing a certain role for sulphur metabolism capability. Several of the genes related to heterotrophy (i.e., ETC complex I), pH stress, low temperature and lipids and biopolymer transporters were acquired through horizontal transfer. Genes related to carotene, high temperature, and salt stress in particular are predicted to have originated at the specific transitioning to the groundwater, rather than inherited from LCA.

The transition of the LCA to soils was marked by higher numbers of genes related to substrate uptake, such as amide, amino acid, lipid or metal ion transport systems in the genome. This likely reflects adaptation to nutrient heterogeneity and patchiness found in terrestrial environments, compared to more homogenous substrates in aquatic environments^71^. Furthermore, stress tolerance mechanisms are more developed compared to other environments including an increased number of genes involved in EPS biosynthesis and export capabilities. This suggests that *Polaromonas* is also able to form biofilms in soils which would allow it to increase nutrient availability and resistance to drying, for instance^72^. We also note the presence of fatty acid desaturases that have been shown to be involved in low temperature stress^73^.

We noted pronounced genomic adaptation when *Polaromona*s transitioned between soils and glaciers. It is plausible that *Polaromonas* transitioned repeatedly between glaciers and proglacial soils during glaciation-deglaciation cycles. In fact, phases of increased dispersal associated with these cycles may have fostered these habitat transitions. Glacier *Polaromonas* genomes obtained genes related to anaerobic respiration (4 genes with 2 transfers), nitrite oxidation (2 genes with 1 transfer), and sulfide oxidation (1 gene), indicating adaptation to oxygen-limited conditions and chemolithotrophic metabolism through horizontal transfer. Furthermore, we note the presence of genes involved in galactose and maltose degradation suggesting specialized carbohydrate metabolism adapted to limited carbon sources. Carbohydrates may originate from snow and ice algae that can form massive blooms in these environments^74^. Interestingly, *Polaromonas* genomes in glaciers lost some genes related to monosaccharides, nitrate and nitrite transport, likely reflecting adaptation to oligotrophic and oxidized conditions in glaciers where such substrates are probably scarcer than in other environments^75^. The presence of sugar degradation genes along with a loss of transporters for the same compounds may indicate that *Polaromonas* in glaciers may favor intracellular metabolisms over the maintenance of multiple transport systems.

In conclusion, our ancestral reconstruction suggests that *Polaromonas*, an iconic and cosmopolitan bacterial genus, radiated from the GFS on the mountain tops downstream into various aquatic and terrestrial ecosystems. The LCA of *Polaromonas* was likely a biofilm former — overall a most successful mode of microbial life — with a versatile metabolism combining heterotrophic and respiratory capacities with broad substrate transport and stress-response systems. These traits reflect adaptation to fluctuating and harsh environmental conditions typical of GFS and glaciers. As *Polaromonas* lineages diversified downstream to colonize aquatic and terrestrial ecosystems, they tended to conserve genes involved in stress response and EPS biosynthesis, suggesting that biofilm formation remains a main trait to adapt to diverse environmental conditions throughout the evolutionary history of this bacterial genus. Our analyses revealed that adaptations through gene gains and losses, as well as horizontal transfers greatly facilitated habitat transitions. Such as acquisition of phototrophic functions and sulfur metabolism in lake and wetland lineages, as well as enhanced nitrogen and lipid uptake systems in groundwater and soils. The distinct adaptations accompanying the various transitions across aquatic and terrestrial ecosystems highlight the genomic plasticity of *Polaromona*s as a key to its success. Our findings also reveal the role of glaciers and streams as a potential cradle for bacterial evolution and diversity.

## Methods

### Microbe Atlas

We used the Microbe Atlas database^39^ to explore the spatial distribution of *Polaromonas* in diverse environments. As *Polaromonas* was found to be a potential contaminant in some kits, we chose a relatively high threshold of abundance to consider it present (> 0.1% relative abundance)^76^.

### Genome gathering

We obtained 632 *Polaromonas* genomes from different sources, including MAGs from the TPMC (160)^43^, TG2G (52)^44^, United Kingdom rivers (8)^19^, GROW DB (4)^18^, GFS (25)^16^ and downloaded via the NCBI dataset software^42^ on the 12^th^ December 2024 (351). The 32 MAGs from Guliya glacier metagenomes^45^ were reconstructed using an ensemble binning and refinement workflow. Contigs were binned with MetaBAT2, MaxBin2, and GroopM2 via UniteM (v0.0.18, https://github.com/donovan-h-parks/UniteM), incorporating sample-specific coverage information from mapped reads, and integrated using UniteM consensus, DAS Tool (v1.1.1)^77^, and metaWRAP (v1.0.6)^78^ bin_refinement. MAG completeness and contamination were assessed using CheckM (v1.0.12), and high-quality bins were further refined by removing GC-and coverage-based outlier contigs with RefineM (v0.0.24; https://github.com/donovan-h-parks/RefineM). Assembly methods are described previously^45^.

We assessed the taxonomy and quality of all MAGs with GTDB-TK (v2.3.0)^46^ and CheckM2 (v1.0.1)^79^, respectively. We then removed genomes that were not annotated as *Polaromonas* (52) and genomes whose completeness and contamination were less than 70% and more than 10%, respectively (248). In order to remove duplicate genomes, we dereplicated them at 99% resulting in a total of 302 genomes using galah (v0.4.2)^80^. As we were interested in only freshwater habitats, we removed strains which were isolated from mine drainage, activated sludge, landfill, permafrost, host-associated and bioreactor environments (20 genomes), with the exception of cultivated species (e.g. *Polaromonas aquatica* CCUG 39402^T^ (GCA 042659145.1), *Polaromonas naphthalenivorans* CJ2^T^ (GCA 000015505.1) and *Polaromonas vacuolata* KCTC 22033^T^ (GCA 012584515.1)) resulting in a total of 282 genomes (Table S1). Interestingly, there were a number of genomes labelled as *Polaromonas* in the NCBI database, which were not in fact *Polaromonas*, but instead being classified as the genus JAAFIP01 or JAAFJR01 in the GTDB. Because of their distinct genomic content, we decided to exclude them from our *Polaromonas* pangenome analysis. We also removed twenty genomes from mine drainage, activated sludge, landfill, host-associated and bioreactor environments to focus on the presence of this genus in natural environments. Due to the way that metadata is deposited in the different databases, we cannot always further detail the type of environments where the samples were taken. Indeed, some soil samples could be linked to permafrost as described elsewhere^13^, while it is usually impossible to find out if glacier samples are from supraglacial, englacial or subglacial parts (Table S1). Further many of the metadata linked to some of the samples were missing such as temperature, salinity concentration or carbon content. Due to recent global datasets being published^16,18,43,44,81,82^, most of the *Polaromonas* publicly available were obtained from samples coming from the Tibetan Plateau (China) (123) followed by Switzerland (29), Canada (25) and the United States (23) (Figure 1a).

### Pangenome analysis

The selected 282 genomes were annotated with Baka (v1.9.3)^83^, microtrait (v1.0.0)^66^ and eggnog-mapper (v2.1.12)^84^. Genomad (v1.11)^62^ was used to identify mobile genetic elements on each genome. Then PPanGGOLiN (v2.1.1)^48^ was used to create the pangenome using default parameters with the exception of the number of partitions defined as 3 (-K). PPanGGOLiN via mmseqs2^85^ (v15) clusters the proteins into gene families with an identity and coverage of 80%. These gene families are then divided into three categories: persistent, shell and cloud. Persistent (1905) and shell genes families (11,722) were then aligned using the “ppanggolin msa” command via the MAFFT software (v7.526)^86^. We analysed the pangenome based on persistent, shell and cloud genes using the occurrence frequencies of genes and their genomic neighbourhood to divide them into persistent, shell and cloud genes that are present in almost all genomes, at intermediate frequencies or low frequencies, respectively^48^.

### Species tree estimation and reconciliation

We then used mrBayes (v3.2.7)^87^ to create gene trees for each gene family using the GTR + Gamma evolutionary model. The Markov Chain Monte Carlo (MCMC) analysis was run for 300,000 generations, with trees sampled every 300 generations and convergence diagnostics assessed every 1,000 generations. We then inferred the species tree using AleRax (v1.0.0)^88^ with the gene trees obtained by mrBayes. An initial species tree was constructed using the MiniNJ (Minimum Evolution Neighbor-Joining) method. To account for missing genes in MAGs, we included a fraction missing file with completeness values for each genome. To reduce runtime, we removed 0.5% of the largest families and we only considered gene families present in at least 30 genomes (total 4,114). The final rooted species tree was inferred using maximum likelihood-based reconciliation approaches via the HYBRID method. Then using this species tree, we performed a reconciliation analysis with the gene trees to determine DTLO (duplications, transfers, losses and origination) parameters which accounts for gene duplication, horizontal gene transfer, and gene loss events. The analysis was performed using a per-family model parameterization, allowing distinct evolutionary models for each gene family and using at 100 sampled gene trees to improve robustness. Ancestral state reconstruction to visualize habitat evolution was done using PastML (v1.9.41)^53^, which calculates ancestral state at each internal node. We used R (v4.5.1)^89^ and the packages targets (v1.11.3)^90^, tidyverse (v2.0.0)^91^, ggtree (v3.16)^92^ and vegan (v2.6-10)^93^ to perform subsequent analyses and plot figures.

## Resource availability

### Lead contact

Further information and requests should be directed to and will be fulfilled by the lead contact, Grégoire Michoud.

### Materials availability

This study did not generate any materials or reagents.

## Data and code availability

The Guliya MAGs are deposited under accession number XXX, while the other MAGs were obtained from public databases. The code to reproduce analyses and figures can be found at https://github.com/michoug/Polaromonas_Pangenome

## Acknowledgments

The Vanishing Glaciers project is supported by The NOMIS Foundation (to T.J.B.). We thank Susheel Bhanu Busi and Daniel Read at the UK Centre for Ecology and Hydrology for their supervision and methodological input into the generation of UK river metagenome-assembled genomes (MAGs), and the UK Environment Agency for supporting sample collection and data generation.

## Author contributions

G.M. was responsible for conceptualization, methodology, software, data curation, investigation, formal analysis, visualization and writing of the original draft. A.G was responsible for conceptualization, methodology, software, data curation, investigation, formal analysis, visualization. H.P. was responsible for conceptualization, methodology, investigation, and writing of the original draft. A.C.T was responsible for data curation. Z.Z was responsible for data curation, investigation. V. R. was responsible for data curation, investigation. T.J.B. was responsible for conceptualization, methodology, investigation, writing of the original draft, supervision, project administration and funding acquisition. All authors commented on previous versions of the manuscript. All authors read and approved the final manuscript.

## Declaration of interests

None

## Notes

### Competing Interest Statement

The authors have declared no competing interest.

