## Supplementary material for "Evolutionary radiation of *Polaromonas* from mountain glaciers downstream": Suplementary data

**Supplementary figures**

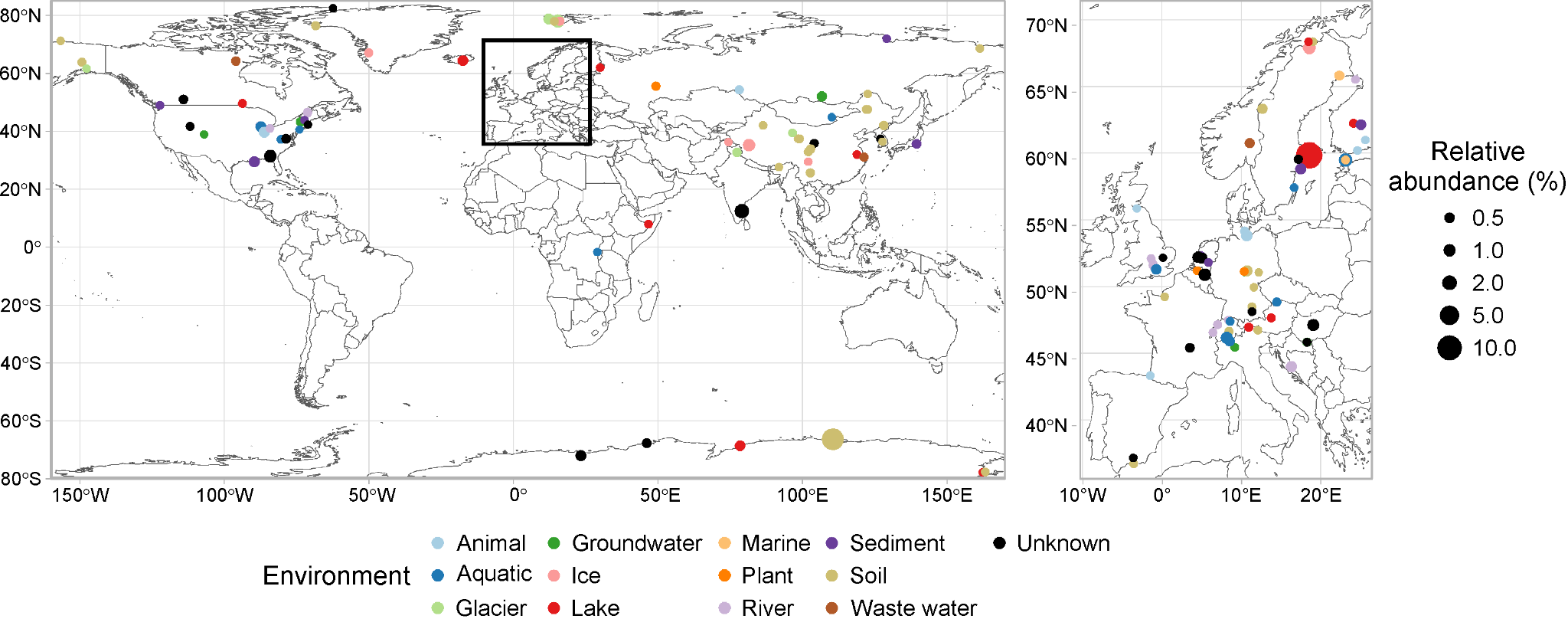

Figure S1: Map representing the geographical locations where members of *Polaromonas* were present above 0.1% of relative abundance as determined by the Microbe atlas project^1^.

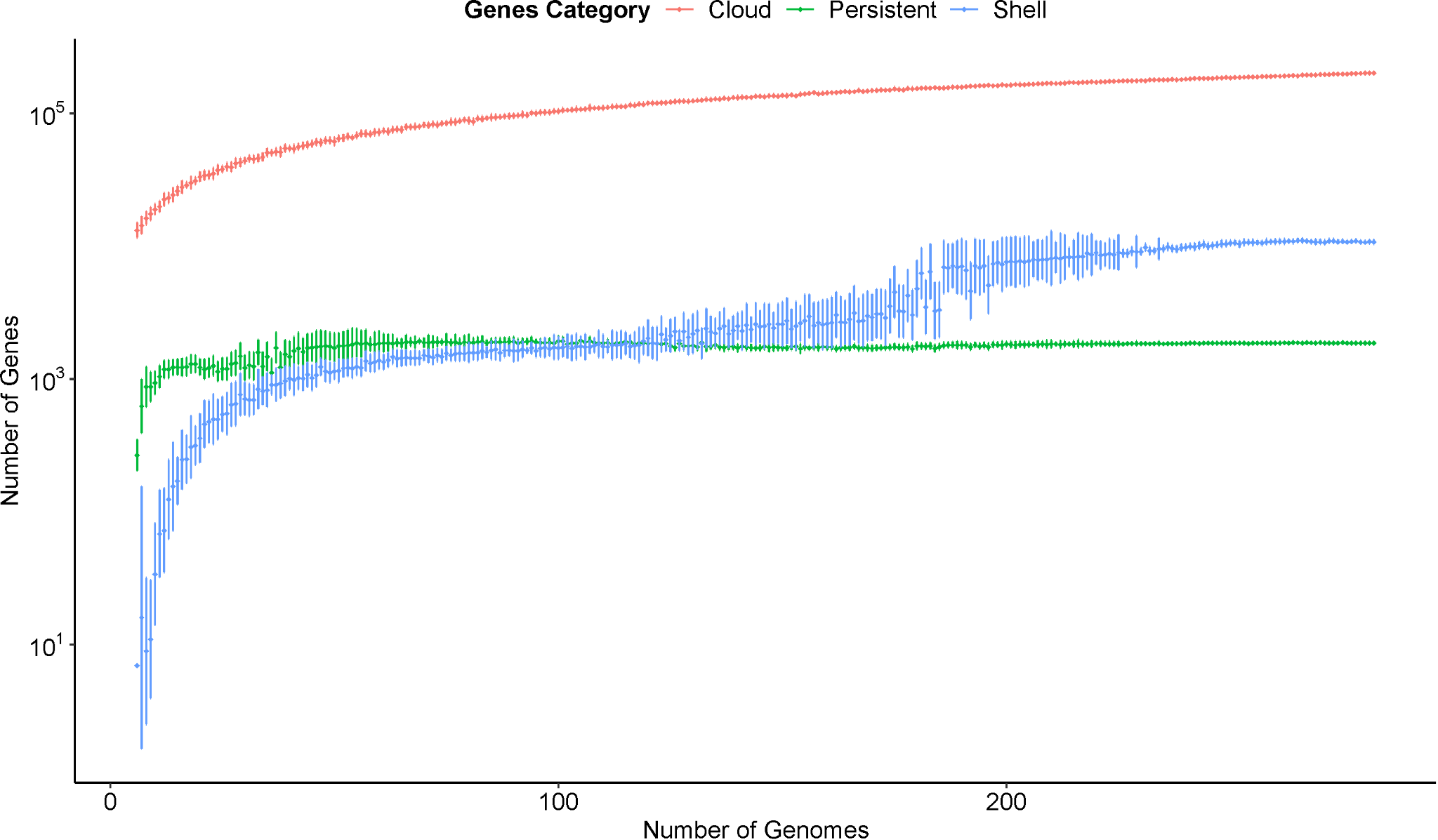

Figure S2: Rarefaction curves of cloud, persistent and shell genes in the *Polaromonas* pangenome.

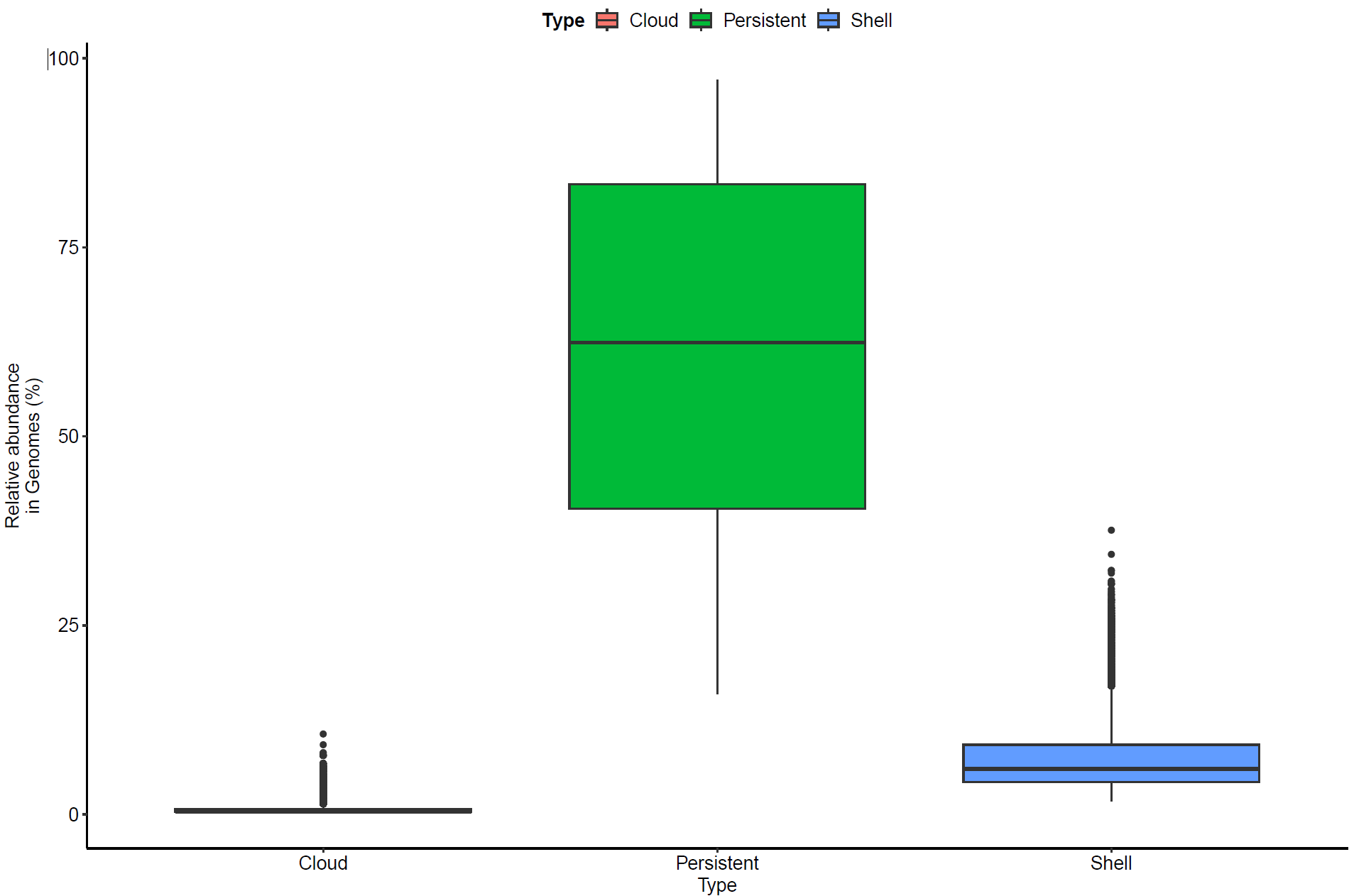

Figure S3: Prevalence of genes classified as cloud, persistent or shell by Ppanggolin^2^ in *Polaromonas* genomes.

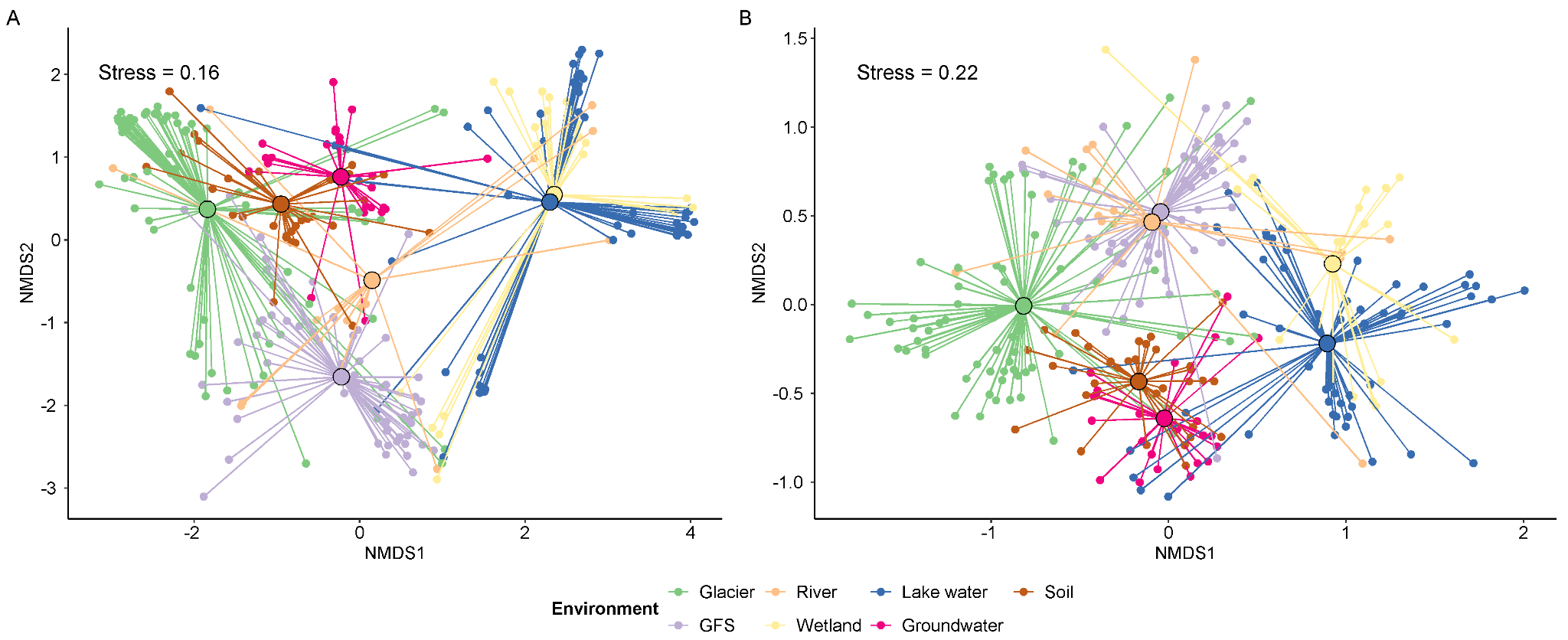

Figure S4: Non-metric multidimensional scaling ordination of shell (A) and cloud (B) genes presence in different genomes show that *Polaromonas* cluster by their environment.

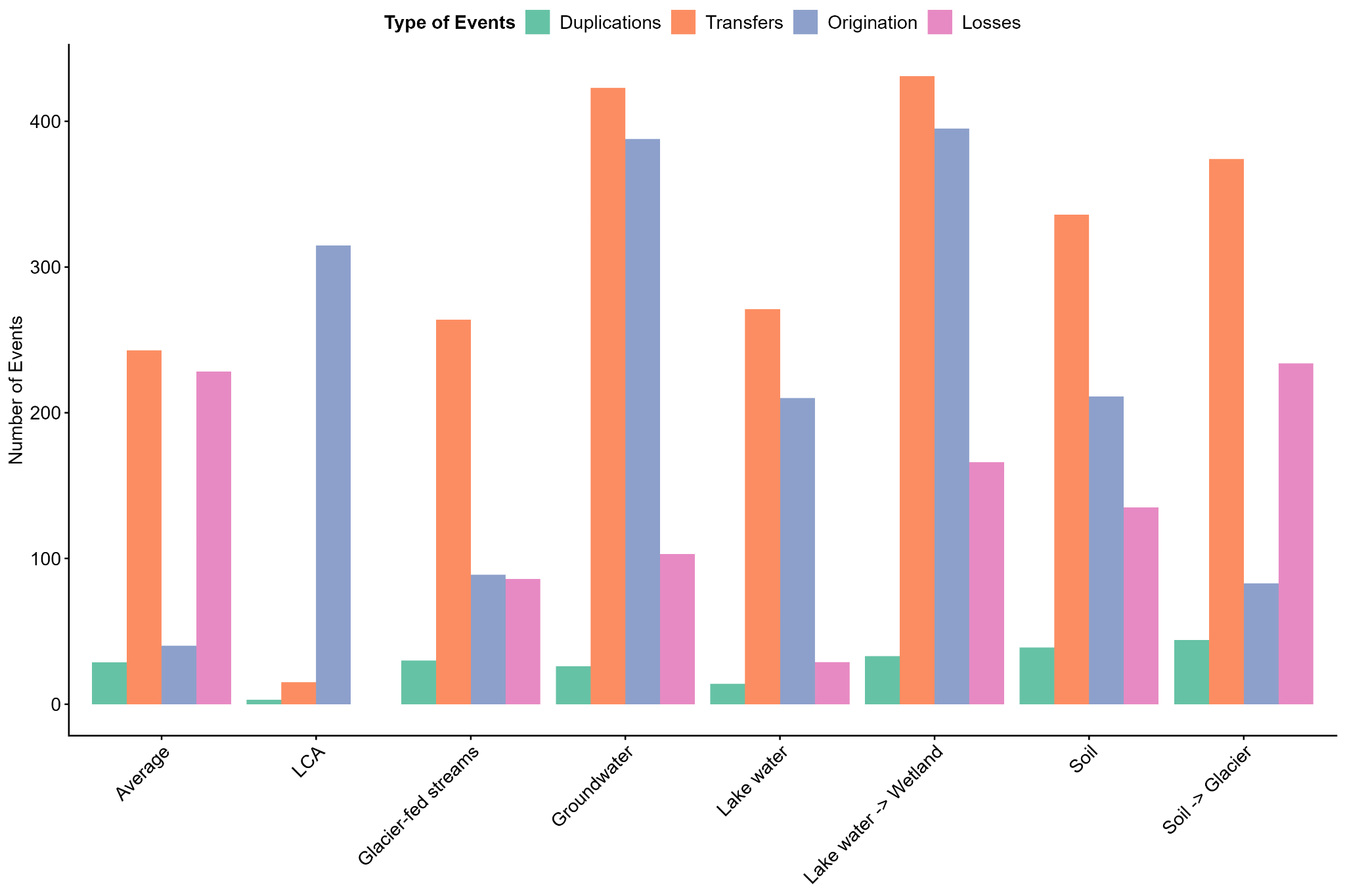

Figure S5: Bar chart showing the number of detected events categorized as duplications, transfers, originations, and losses across different habitat transitions (LCA, Glacier, Glacier-fed streams, Wetland, Lake water, Groundwater, and Soil). The “Average” column represents the mean number of events across all internal nodes.

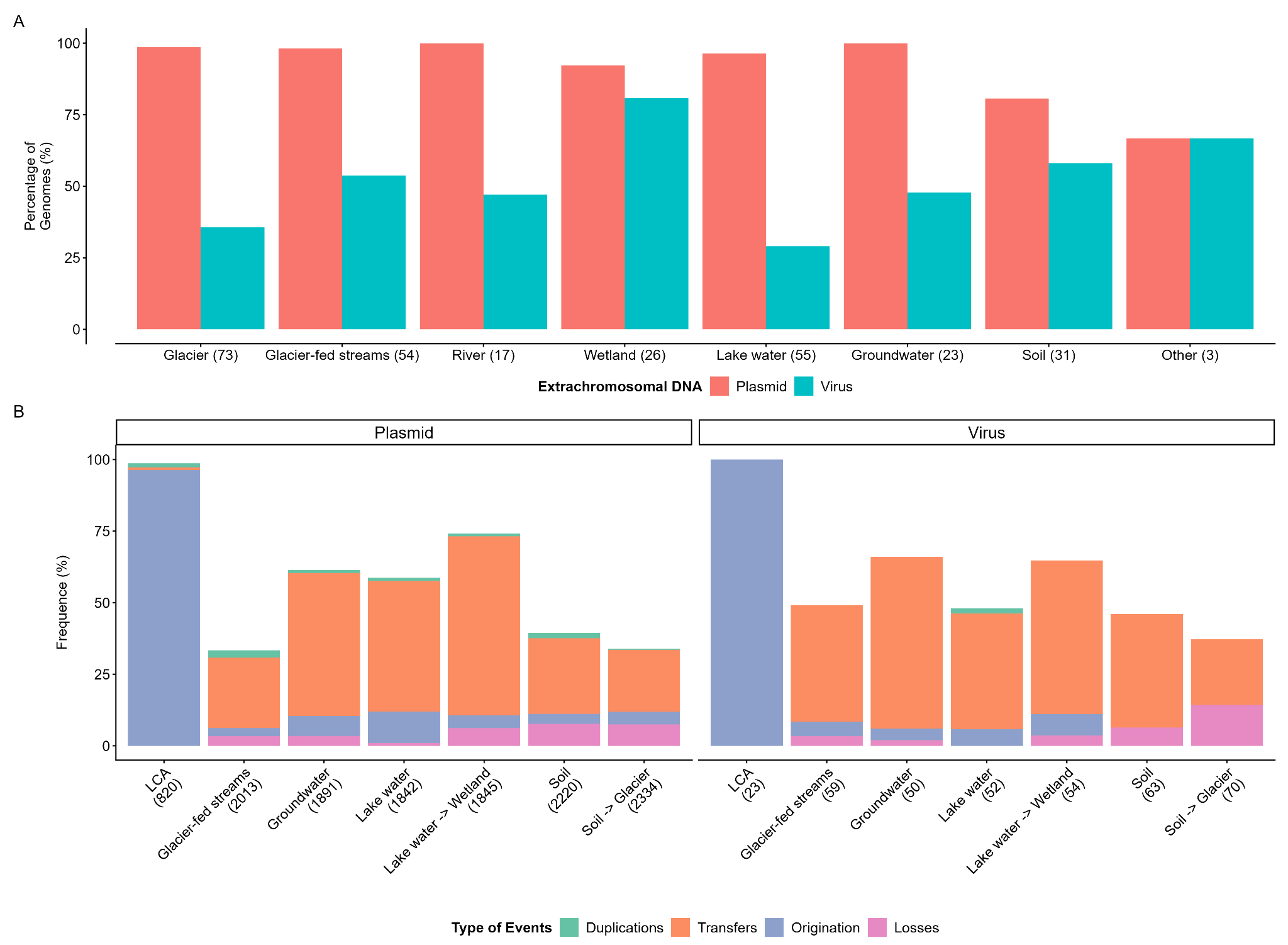

Figure S6: (A) Presence of predicted viruses and plasmids associated with the different *Polaromonas* genomes. The prediction of such mobile genetic elements was obtained using genomad^3^. Numbers associated with each environment correspond to the number of genomes belonging to such biomes. (B) Frequency of mobile genetic elements found at habitat transitions. Colors correspond to the predicted events that occurred at which transitions. The remaining uncolored part represents mobile genetic elements vertically transmitted (via normal bacterial division).

Table S2: Characteristics of the *Poloramonas* genomes studied grouped by environments. The relative length was determined by normalizing the observed length by the completeness.

| Environment | Number of genomes | Relative length (Mbp) | GC (%) | Completeness (%) | Contamination (%) |
| --- | --- | --- | --- | --- | --- |
| Glacier | 73 | 3.99 ± 0.47 | 60.2 ± 2.4 | 90.2 ± 10.0 | 0.7 ± 0.8 |
| Glacier-fed streams | 54 | 4.00 ± 0.53 | 58.3 ± 2.1 | 82.8 ± 8.0 | 2.3 ± 1.9 |
| Groundwater | 23 | 4.28 ± 0.58 | 62.5 ± 1.6 | 90.6 ± 7.1 | 2.9 ± 2.7 |
| Lake water | 55 | 3.74 ± 0.72 | 60.9 ± 2.2 | 83.7 ± 10.0 | 1.1 ± 1.3 |
| Other | 3 | 4.97 ± 1.00 | 58.1 ± 5.2 | 99.8 ± 0.4 | 2.3 ± 2.3 |
| River | 17 | 3.74 ± 0.53 | 60.9 ± 2.2 | 83.5 ± 8.6 | 1.3 ± 1.3 |
| Soil | 31 | 5.09 ± 1.26 | 62.0 ± 1.5 | 87.7 ± 11.5 | 1.6 ± 2.4 |
| Wetland | 26 | 3.51 ± 0.49 | 61.0 ± 2.5 | 80.6 ± 6.2 | 1.1 ± 1.2 |
| All | 283 | 4.04 ± 0.80 | 60.4 ± 2.5 | 86.1 ± 9.9 | 1.4 ± 1.8 |
